# ApoE4 attenuates cortical neuronal activity and impairs memory in young apoE4 rats

**DOI:** 10.1101/2021.01.14.426651

**Authors:** Ilona Har-Paz, Elor Arieli, Anan Moran

**Affiliations:** Department of Neurobiology, The George S. Wise Faculty of Life Sciences, Tel Aviv University, Tel Aviv, Israel 6997801; Sagol School of Neuroscience, Tel Aviv University, Tel Aviv, Israel 6997801

**Keywords:** ApoE4, Neuronal activity, Taste learning, Alzheimer’s disease, Neuronal information coding

## Abstract

The E4 allele of apolipoprotein E (apoE4) is the strongest genetic risk factor for late-onset Alzheimer’s disease (AD). However, apoE4 may cause innate brain abnormalities before the appearance of AD related neuropathology. Understanding these primary dysfunctions is vital for early detection of AD and the development of therapeutic strategies for it. Recently we have shown impaired extra-hippocampal memory in young apoE4 mice – a deficit that was correlated with attenuated structural pre-synaptic plasticity in cortical and subcortical regions. Here we test the hypothesis that these early structural deficits impact learning *via* changes in basal and stimuli evoked neuronal activity. We recorded extracellular neuronal activity from the gustatory cortex (GC) of three-month-old humanized apoE4 and wildtype rats, before and after conditioned taste aversion (CTA) training. Despite normal sucrose drinking behavior before CTA, young apoE4 rats showed impaired CTA learning, consistent with our previous results in apoE4 mice. This behavioral deficit was correlated with decreased basal and taste-evoked firing rates in both putative excitatory and inhibitory GC neurons. Single neuron and ensemble analyses of taste coding demonstrated that apoE4 neurons could be used to correctly classify tastes, but were unable to undergo plasticity to support learning. Our results suggest that apoE4 impacts brain excitability and plasticity early in life and may act as an initiator for later AD pathologies.

**Significant statement:** The ApoE4 allele is the strongest genetic risk-factor for late-onset Alzheimer’s disease (AD), yet the link between apoE4 and AD is still unclear. Recent molecular and in-vitro studies suggest that apoE4 interferes with normal brain functions decades before the development of its related AD neuropathology. Here we recorded the activity of cortical neurons from young apoE4 rats during extra-hippocampal learning to study early apoE4 neuronal activity abnormalities, and their effects over coding capacities. We show that apoE4 drastically reduces basal and stimuli-evoked cortical activity in both excitatory and inhibitory neurons. The apoE4-induced activity attenuation did not prevent coding of stimuli identity and valence, but impaired capacity to undergo activity changes to support learning. Our findings support the hypothesis that apoE4 interfere with normal neuronal plasticity early in life; a deficit that may lead to late-onset AD development.

## Introduction

The prevalence of the E4 allele of apolipoprotein E (apoE4) gene in late-onset Alzheimer’s disease (AD) patients is well established (Huang, 2011; Lane-Donovan and Herz, 2017). The observed AD-apoE4 link has been attributed to an increased amyloid-beta (Aβ) accumulation and tau phosphorylation *via* lipid-related mechanisms (Huang, 2011; Verghese et al., 2013; Abushakra et al., 2016; Zhao et al., 2018; Fernandez et al., 2019). However, evidence suggests that these pathologies are secondary to earlier basal anomalies that interfere with the structure and function of normal brains, including simpler dendritic structures and disrupted plasticity and clearance mechanisms (Baxter et al., 2003; M. Di Battista et al., 2016).

Most of the work on these primary and secondary apoE4 deficits has focused on the hippocampus, where they were observed simultaneously at early ages (Trommer et al., 2004; Chen et al., 2010; Yin et al., 2011; Leung et al., 2012; O’Dwyer et al., 2012; Liraz et al., 2013). This may make it difficult to identify whether and how apoE4 primary anomalies affect the development of the later deficits. In contrary, cortical morphological deficits in neurons of layers II/III (Dumanis et al., 2009) and amygdalar reduced basal excitatory transmission (Wang et al., 2005) have been identified in young apoE4 mice in the absence of AD neuropathology. Investigating these extra-hippocampal systems during early ages may therefore provide valuable insight into the pre-pathological effects of apoE4 on brain functions. We recently showed structural plasticity impairments in intact cortical and subcortical regions to be correlated with impaired learning of a cortically and amygdalar-dependent but hippocampal-independent form of learning called conditioned taste aversion (CTA, Har-Paz, Roisman, Michaelson, & Moran, 2019) in target-replacement (TR) apoE4 mice. ApoE4, therefore, seems to reduce plasticity even in younger animals and extra-hippocampal brain tissues.

Neuronal activity drives plasticity, which in turn changes neuronal activity. Therefore, the apoE4-related reduced structural plasticity may impact neuronal activity affecting functional plasticity deficits. We hypothesized that the extra-hippocampal spiking activity may be reduced similarly to findings from the amygdala in slice preparations (Wang et al., 2005). Moreover, we also aimed to test whether apoE4 impacts the capacity of neurons to code information and drive coding related adaptations following learning. To test these hypotheses, we compared the basal activity and taste responses of single neurons in the GC between young (3-month-old) knocked-in humanized apoE4 (hApoE4) and wildtype (WT) rats, before and after CTA learning. The GC is uniquely suited for this study as its neuronal activity has been robustly characterized in rats. Specifically, GC neurons code taste identity and palatability in distinct early (1st second post taste stimulation) and late (1-2 seconds post taste stimulation) response epochs (Katz et al., 2001; Piette et al., 2012; Sadacca et al., 2012). This epoch-specific coding is modified by learning, both at the single neuron and ensemble levels, in a specific manner that reflects changes in the hedonic value of a taste (Grossman et al., 2008; Moran and Katz, 2014). These response properties provide a rich basis upon which to portray the impact of apoE4 on the capacity of GC neurons to encode and transmit taste information.

The behavioral results recapitulated our previous findings using TR apoE4 and apoE3 mice, showing impaired CTA learning in the hApoE4 compared to WT rats. Furthermore, our basic hypothesis was confirmed in that cortical neurons of hApoE4 rats fired at a much lower frequency compared to WT, an effect that was particularly large in putative inhibitory interneurons. Also, we found that although apoE4 GC neurons could classify taste information correctly (both identity and palatability) using their reduced firing rates, they fail to appropriately update their palatability coding following the aversive conditioning. We discuss our results in relation to the hypothesis of inherent apoE4 vesicular flux and plasticity deficits that can contribute to the development of AD pathology in aged carriers.

## Methods

### Subjects

Young adult (3-month-old, 250-300 gr) Sprague Dawley (SD) female humanized apoE4 (hApoE4, n=13), and wildtype (WT, n=10) rats were utilized in this study. Animals were subjected to a 12 hours dark/light schedule with the experiments performed in the light portion of the cycle. Rats were given ad libitum access to chow and water unless otherwise specified. All methods complied with the Tel-Aviv university Institutional Animal Care and Use Committee guidelines.

### Surgery

Rats were anesthetized using an intraperitoneal injection of a ketamine/xylazine mixture (100 and 10 mg/kg, respectively; maintenance: one-third induction dose every 1h), and placed into a stereotactic frame. The scalp was excised and holes were drilled in the skull for the insertion of 4 self-tapping ground screws and a bundle of 32 formvar-coated, 25 μm nichrome wire bundle (AM-Systems) attached to a mini-microdrive (Moran and Katz, 2014) 0.5 mm above the GC (AP=1.4 mm ML=5 mm from bregma; DV=4.5 mm from dura). Additionally, an intraoral cannula (IOC) was inserted between the first maxillary molar and the lip, exiting close to the skull ridge (Phillips and Norgren, 1970; Katz et al., 2001). The electrode connector and the IOC were then secured to place using dental cement. The rats were given a week to recover before experiment initiation.

### Experimental Setup

The experimental cage (Coulbourn Instruments) containing two infrared (IR)-operated nose-pokes resides in a sound attenuation box that included a Faraday cage to reduce electromagnetic interference. The experiment was controlled by an Arduino based system (Arduino.cc, 2015) that responded with a delivery of a 40μL drop of liquid directly into the oral cavity of the rat upon breakage of the nose-pokes’ infrared beam. A timestamp (in milliseconds) of every such event was recorded by an Intan recording system (Intan Technologies) that was also responsible to record the signals from the 32 electrodes (30,000 samples/second per channel) implanted in the GC during the entire session.

### Experimental procedure

The rats were handled 15 minutes per day for 3 consecutive days to reduce stress before the experiment. During the experiment, the rats were habituated to a scheduled water regime in which access to fluids was allowed for 15 minutes in the morning recording session, and 2 hours in the afternoon (to allow proper hydration). Each morning recording session was divided into two parts: an initial behavior assessment followed by a neuronal response session (Fig. 1). In the behavior assessment part, the rats were allowed to poke in two nose-pokes to receive 40μL drops of fluids via the IOC for 15 minutes (minimum of 3 seconds inter-taste interval). In the second part, pseudo-randomly selected deliveries (with equal probability) of different tastes (12 seconds inter-taste interval, 30 trials each) were delivered through the IOC. This enabled the recording of the GC electrophysiological signals alongside the exact timestamps of stimuli deliveries. During the initial 3 habituation days water was given in the first part and 0.1M NaCl, water or 0.1M citric acid (CA) in the second part. On the 4th day (the training session), novel 0.2M sucrose was given in the first part and was also added to the battery of tastes delivered in the passive delivery part. This session was followed by an IP injection of LiCl (0.15M, at 1% body weight) for the induction of gastrointestinal malaise. The next day we tested for sucrose aversion by allowing the rats to choose between two nose-pokes, one that administered drops of water and the other sucrose. The passive taste delivery protocol was then conducted similarly to the previous session.

**Figure 1:**
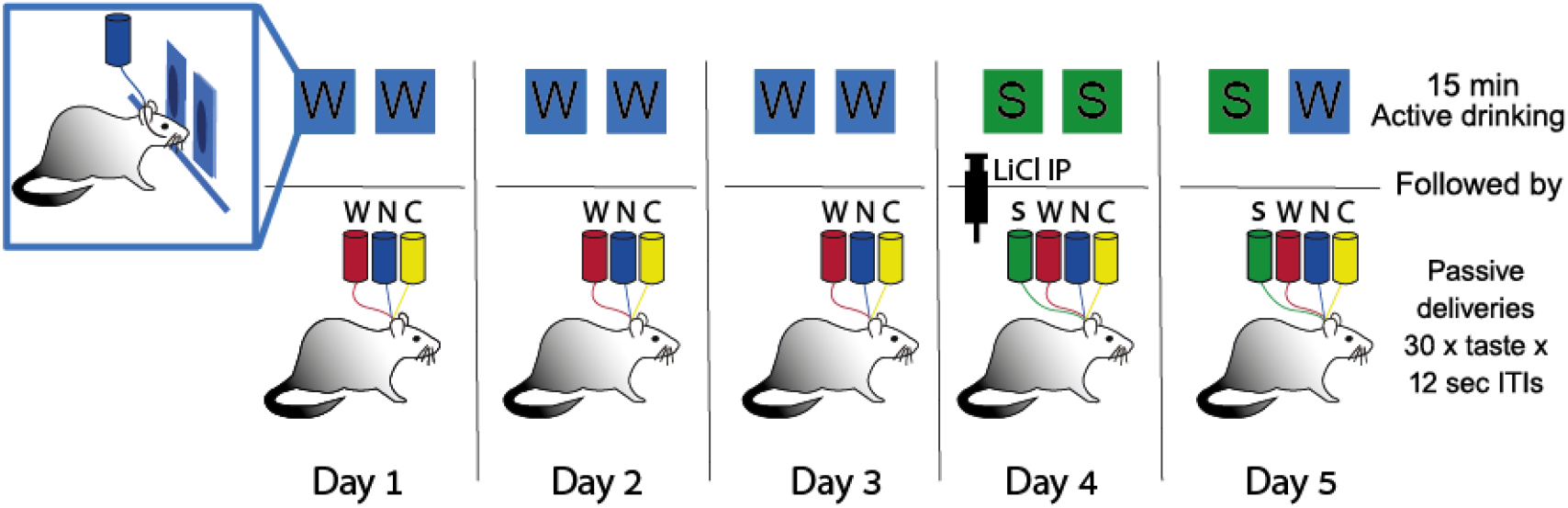
Experimental protocol. The behavioral protocol started with 3 habituation days in which the rats learn to poke in two nose-pokes to receive drops of water via the IOC. This was followed by passive pseudo-random taste deliveries of NaCl (N), water (W) and citric acid (C) solutions that were provided every 12 seconds, 30 trials for each taste. On day 4 (“Training day”) a novel sucrose solution (S) was emitted upon activation of both nose-pokes, and was also added to the subsequent passive taste deliveries. This session was followed by a LiCl injection to evoke gastrointestinal malaise. On the next day CTA level was assessed by calculating the ratio between poking for sucrose and water operated by two different nose-pokes in the experimental chamber. Neural activity was recorded during the passive deliveries part of the training and testing days. ITI = inter-taste interval.

### Exclusion criteria

Rats were excluded from the experiment if one of the following criteria were met: Over 20% weight reduction from their pre-surgery weight, apparent illness or inflammation signs, behavior which indicates un-wellness (e.g., lack of drinking during the experimental session).

### Poking bout analysis

The two nose-pokes placed in the experimental cage allowed for liquid deliveries into the rat’s oral cavity as a result of a poking activity in 3 seconds intervals. The rats’ poking patterns analysis (Arieli et al., 2020) was adapted from previous licking analysis protocols (Baird et al., 2005; Neath et al., 2010; Lin et al., 2012) to assess differences in sucrose palatability between the genotypes. Accordingly, a drinking bout was defined as a consecutive drinking period with intervals smaller than 5 seconds between one delivery to another. An inter-bout interval was defined as a drinking break with a minimal span of 5 seconds.

### Sucrose detection test

Differences in sucrose concentration perception between the genotypes were tested using consecutive two-bottle tests: Single-housed rats were presented in their home cage with one bottle containing water and another bottle containing a certain concentration of sucrose solution daily for 24 hours. Every 24 hours the bottles were replaced and randomly placed in the cages. The new sucrose bottle always contained half the concentration of the previous day (0.1M, 0.05M, 0.025M, 0.0122M, 0.006M and 0.003M). The amounts drank from each bottle were measured and sucrose preference (sucrose intake divided by the total intake) was calculated as a measure of assessing sucrose detection by the rats (using rodents’ innate tendency to choose sucrose over water if detected).

### Spike detection and spike sorting

Spike sorting was performed using the open-source KlustaKwik/Phy software (Rossant et al., 2015), and conducted on each electrode signal at a time. The electrophysiological signal collected from each electrode was first high-pass filtered (350Hz cutoff) to eliminate slow fluctuations, and then a moving threshold of 3-6 standard deviations was calculated to identify putative spikes. These spikes were first automatically clustered by the Klusta-Kwik algorithm, and refined manually using the Phy graphical user interface to produce well-isolated units. Only stable units that showed a clear refractory period (1 ms) were included in this study.

### Single unit analysis

All data analyses were done using self-made scripts written in Python and available in the GitHub repository (https://github.com/ananmoranlab/moran_lab). After obtaining the final single units from each group (apoE4 pre-CTA, apoE4 post-CTA, WT pre-CTA, WT post-CTA) firing rates (FRs) were calculated for each unit using 50 milliseconds which were then averaged into a peri-stimulus time histogram (PSTH, from ~120 trials, which were aligned to each of the taste stimuli containing 30 trials per tastant) and smoothened using a Savitsky-Golay filter for visualization. The baseline (BL) FR was calculated from the one second that preceded the taste deliveries. Single unit evoked taste responses were normalized to their averaged BL before averaging per group. Normalized taste responses were analyzed either over the span of 3 seconds after taste delivery, during the early epoch (EE, 0-1 second after taste delivery), and the late epoch (LE, 1-2 second after taste delivery).

### Neuron type classification

Classification of GC neurons as either putative excitatory or inhibitory was based on two parameters of a spike shape: the width (W) and height (H) of the spike (Mitchell et al., 2007; Yokota et al., 2011; Samuelsen et al., 2012). Specifically, random 10% of each unit’s spikes were selected and aligned to its peak and averaged to produce an exemplary spike shape of the neuron. For each averaged waveform we then calculated the ratio (H/W) between the height (measured from BL to peak) and width (measured at half the height). The H/W ratio values over all neurons showed a clear bimodal distribution (Fig. 3A). Ensuing unsupervised classification of the H/W values using the K-means algorithm with 2 clusters produced 2 groups of neurons: one with a narrow spike shape (defines as putative inhibitory neurons), and the other with a wider shape (defined as putative excitatory) (Fig. 3A inset) Further confirmation of the validity of our method came from FR analysis showing significantly higher firing rate of the inhibitory group compared with the excitatory group (Mitchell et al., 2007; Yokota et al., 2011; Samuelsen et al., 2012).

### Taste responsivity and specificity characterization

A neuron was classified as taste responsive if its average FR during a 2-second window post taste delivery was either 3 standard deviations over or under its baseline FR. Taste specific neurons were identified using Two-way ANOVA over time (0.25s bins post stimuli delivery) and taste (water, NaCl, CA, sucrose). A neuron was defined as “taste specific” if the ANOVA’s main factor of taste or the time-taste interaction were significant with p<0.0001.

### Palatability index (PI) calculations

We evaluated a neuron’s PI in the time following taste stimulation by calculating the correlation between its taste responses and the intrinsic averseness/agreeability of the tastes (Piette et al., 2012; Sadacca et al., 2012). Initially, the neuronal taste responses were binned in 50ms bins and averaged over trials of the same tastant into an averaged taste post-stimulus time histogram (PSTH). We then calculated the bin-wise squared correlation between the PSTH and the known tastes’ palatability. The resulted correlation values were then used to test differences in basal PI between the apoE4 and WT genotypes.

### Single neuron palatability classifications

The responses for each of the four tested stimuli (NaCl, water, sucrose, and CA) were averaged separately for each single neuron over 250 ms time bins in the 3 s window after each taste delivery (30 trials per taste). This created 4 PSTHs per neuron. Each neuronal single response for a taste was binned similarly (250 ms bins over 3 s window) and the Euclidean distance was then calculated between the single trial and each of the PSTHs. A taste response in a single trial was classified as evoked by a certain tastant if its Euclidean distance from that stimuli PSTH was the smallest.

### Evaluating sucrose palatability based on population response dynamics

GC Population-level sucrose PC was based on calculating the relative distance between sucrose ensemble responses to either palatable NaCl or aversive CA. Using a 3D matrix of trials × neurons × bins for each taste we first averaged over trials to get the 2D mean population vector for each bin in the response dynamics, and then reduced the dimensionality of the neurons dimension to 2 using principal component analysis (PCA). All taste responses of each rat separately were calculated and reduced together and then plotted on the same PCA space to allow for comparisons. We then calculated the average Euclidean distance in the PCA space at certain windows (EE or LE) of the response dynamics, and defined sucrose palatably coding (PC) as the ratio between CA-sugar and NaCl-sugar distances. A high PC ratio indicates that the population response dynamics to sucrose resembled NaCl rather than CA in the coding space, thus indicating positive sucrose PC.

### Statistical analyses

The results in the graphs are expressed as mean ± SEM. Two-way ANOVA was performed to compare the FR and neuronal differences between the groups unless otherwise specified. For differences in post-CTA drinking behavior, we utilized independent samples welch corrected t-test. Differences between the neuronal population properties recorded in the different groups were analyzed using the chi-square test. Tukey-corrected multiple comparisons test was performed to assess within-group changes. The value in parenthesis of the F, t, and X^2^ statistics represent degrees of freedom. The significance level was set to 0.05 for all statistical analyses.

## Results

Young adult humanized ApoE4 (hApoE4) and wildtype (WT) SD rats were used in this study. Rats were trained in a conditioned taste aversion (CTA) paradigm: entering an IR-armed nose poke caused sucrose to be delivered via an intraoral cannula (IOC) directly into the oral cavity of the rats for 15 minutes (Fig. 1). Ensuing deliveries of tastes though the IOC, simultaneously with GC neuronal recording, provided an assessment of neuronal responses. An IP injection of the emetic LiCl was given at the end of this session to induce malaise (Fig. 1) (Arieli et al., 2020). Recently we showed that CTA learning is impaired in young TR apoE4 mice (Har-Paz et al., 2019). To study the generality of this phenotype we tested whether a similar memory impairment is evident in young adult hApoE4 rats. Three months old WT (n=9) and hApoE4 (n=9) rats consumed similar amounts of water on the last habituation day (t-test: t(16)=0.43, p=0.67), and sucrose solution on the training day (Fig. 2A, t-test: t(16)=0.33, p=0.73).

**Figure 2.**
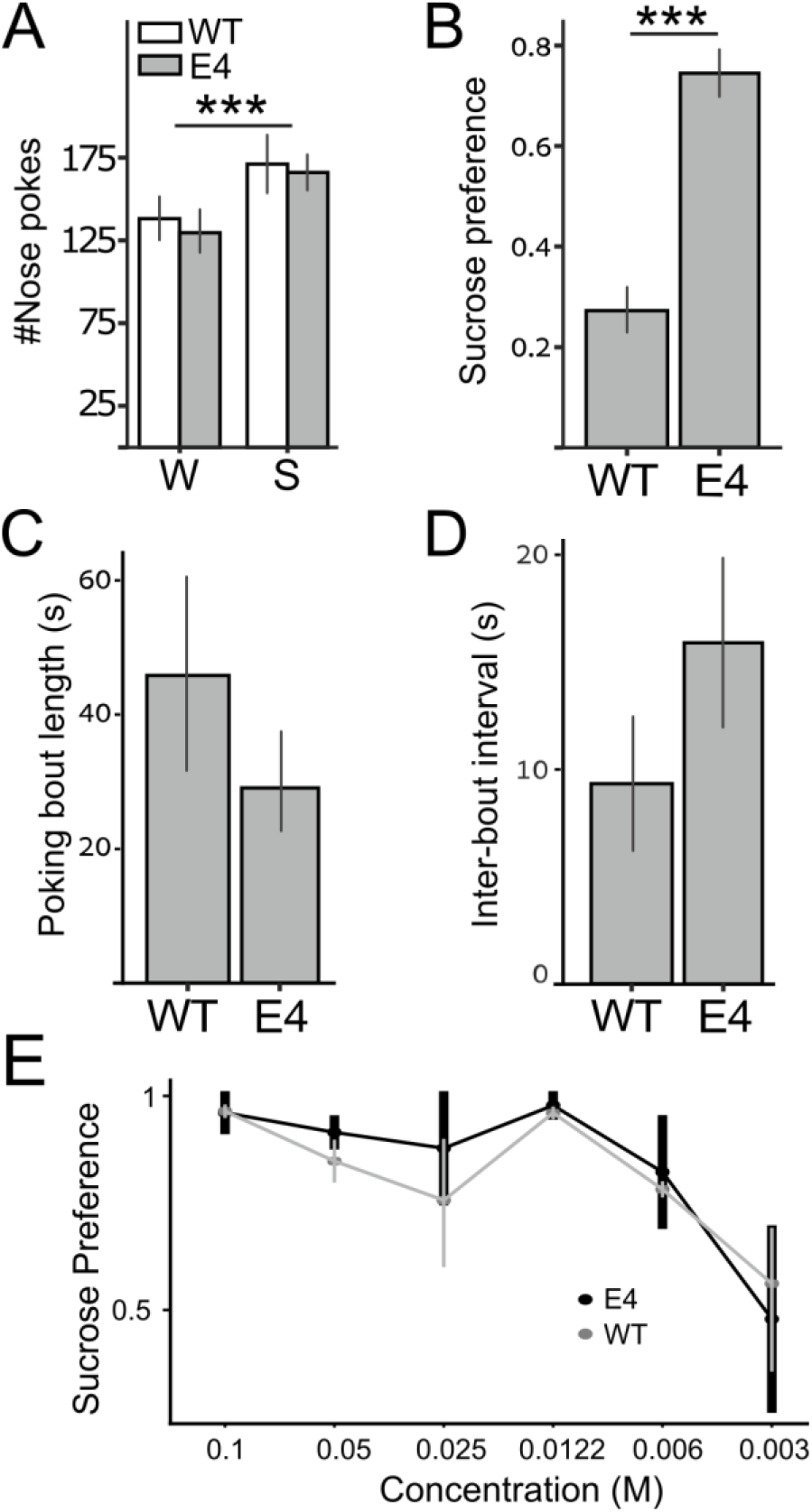
Impaired CTA learning in apoE4 rats. **A)** Sucrose (S) intake levels on test day exceeded those of water (W) on the last habituation trial with no genotype related differences. **B)** ApoE4 rats show impaired CTA memory on test day as manifested by their high sucrose preference compared to WT. **C-D)** Licking and poking dynamics are used to infer taste palatability. No significant genotype differences were found in poking bouts length. **(C)** and inter-bout intervals **(D)** to sucrose during the training session, indicating similar palatability perception. **E)** Sucrose preference was measured at different sucrose concentrations revealing no significant differences between the genotypes. Sucrose preference = Sucrose Intake out of total water and sucrose intake; E4 = ApoE4; WT = Wildtype; **\***\*\* p<0.001

The two groups preferred sucrose similarly prior to training compared to water (Fig. 2A, Two-way ANOVA, genotype: F(1,31)=0.274, p=0.603; taste: F(1,31)=4.164, p=0.049; Interaction: F(1,31)<0.0001, p=0.99). Twenty-four hours after training, however, when the level of CTA learning was assessed in a comparison of sucrose intake to that of water during a test session (Fig. 1, Day 5), a significant difference between the groups appeared: the WT group showed evidence of a strong CTA by avoiding the sucrose nose-poke, while the apoE4 rats continued to consume significantly more sucrose than water (Fig. 2B, t(16)=6.74, p<0.001). An additional set of tests further confirmed that the impaired CTA learning is not due to group differences in palatability perception: A poke-pattern analysis revealed similar poking bout lengths (Fig. 2C, t(16)=0.34, p=0.74) and inter-bout intervals (Fig. 2D, t(16)=0.62, p=0.55) between the groups, an indication that palatability perception in the apoE4 rats was normal (Arieli et al., 2020). Additionally, a test comparing preferences for different sucrose concentrations showed only the expected preferences for higher concentrations and no between-group differences (Fig. 2E, Repeated measures Two-way ANOVA, genotype: F(1,5)=0.504, p=1; sucrose concentration: F(5,25)=5.21, p=0.002; Interaction: F(5,25)=0.21, p=1). Overall, the current results confirm our previous findings in mice (Har-Paz et al., 2019), demonstrating apoE4-related impaired CTA learning in young adult animals, despite intact taste perception. This remarkable agreement across species further supports the use of the CTA paradigm in the study of the neuronal activity abnormalities in young apoE4 animals.

Neuronal activity in the GC is involved in taste perception (Katz et al., 2001; Piette et al., 2012; Sadacca et al., 2012) and CTA is known to be subserved by changes of this activity (Yasoshima and Yamamoto, 1998; Moran and Katz, 2014). The impaired CTA in apoE4 rats, therefore, may reflect normally active neurons that only lack sufficient synaptic resources to support the required plasticity during learning (Har-Paz et al., 2019). However, it is possible that apoE4 also alters different basal properties of the neurons, and therefore impacts basic taste processing. If so, we assumed that these alterations can be detected in the baseline (BL) and neuronal taste responses prior to learning. To evaluate these options, we conducted electrophysiological recordings in the GC of awake apoE4 and WT rats, before and 24 hours after CTA induction. Neuronal activity was measured in the sessions that immediately followed behavioral testing, in which rats received samples of multiple tastants through the IOC (Fig. 1). The pre- and post-CTA BL-FR histograms of the two genotypes show a typical exponent distribution (Fig. 3A), but the apoE4 group shows higher occurrence of low firing neurons. Across all neurons, the averaged BL-FR of neurons from the hApoE4 rats was significantly lower than the WT rats both before (Fig. 3B left, Tukey: WT Pre vs. apoE4 Pre p=0.001), and after (Fig. 3B right, Tukey: WT Post vs. apoE4 Post p=0.001) CTA. This lower BL activity suggests the existence of abnormalities in the innate excitability of GC neurons in young apoE4 animals.

**Figure 3.**
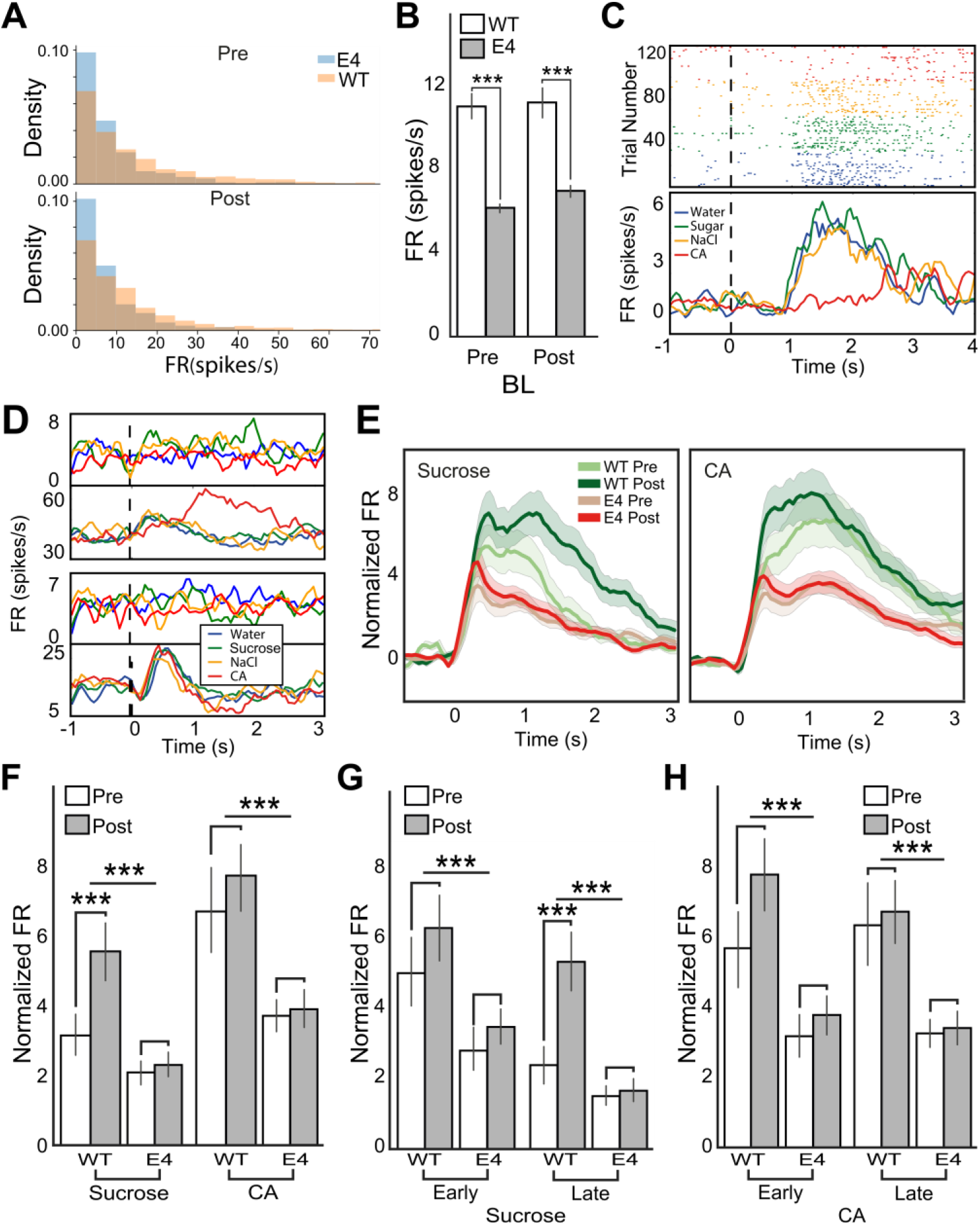
Reduced neuronal firing rate and plasticity in the GC of apoE4 animals. **A)** The distribution of basal activity in GC neurons. More neurons of the apoE4 group show low FR compared to WT. **B)** The average BL FR of apoE4 GC neurons is significantly lower than those from the WT group. **C)** Spike-time rasters (top) and PSTHs (bottom) for an examples of GC neuronal responses to water, NaCl, sucrose, and CA from a WT rat. **D)** A collection of GC neuronal responses from the apoE4 (top two) and WT (bottom two) rats. **E)** Averaged normalized taste responses over all neurons for each genotype and day groups; left panel-sucrose, right panel – CA. **F)** Mean GC neuronal response amplitude over the 3 seconds after taste delivery (left – sucrose, right – CA), pre- and post-CTA. Learning increased WT responses to sucrose, but not in the apoE4 group. **G)** Sucrose neural response divided into epochs. A strong genotype effect was evident in both epochs, while the LE revealed a specific post-CTA FR increase only in WT animals. **H)** Similar as in (H) but for CA, showing only the genotype effect without day (learning) effect. FR = firing rate; Normalized FR = the FR of each neuron normalized to its baseline activity; Early, EE = 0-1 second post taste delivery; Late, LE = 1-2 second post taste delivery; WT = Wildtype; E4 = apoE4; Pre = before CTA learning during the training session; Post = after CTA learning during the test session; *** p<0.001.

These lower BL levels of activity might be expected to impact taste response patterns, and to interfere with normal taste processing. In normal rodent brains, many GC neurons respond dynamically to taste stimuli, changing their FR patterns multiple times across several seconds (Fig. 3C, Katz et al., 2001; Piette et al., 2012; Sadacca et al., 2012). Initial visual inspection of GC neural responses to tastes did not reveal clear differences between apoE4 (Fig. 3D, upper 2 panels) and WT animals (Fig. 3D, lower 2 panels)—neurons from both groups displayed a plethora of dynamic responses, and similar fractions of the populations were taste-responsive (Table 1, χ^2^-test, χ^2^(4, N=624)=6.84, p=0.07).

**Table 1:** A summary of the neuronal populations recorded in this study for each genotype either before CTA learning (Pre) or after CTA learning (Post), E4=ApoE4 rats, WT=wildtype rat; Taste responsive = significant response to one or more of the tastants compared to the BL activity; Taste specific = responded in a distinct manner to at least one tastant.

|  | E4 |  | WT |  |
| --- | --- | --- | --- | --- |
|  | Pre | Post | Pre | Post |
| **Taste responsive** | (80.6%) 179 | (84.4%) 157 | (83.7%) 72 | (83.7%) 96 |
| **Taste specific** | (10.3%) 23 | (20.9%) 39 | (33.6%) 22 | (33.6%) 39 |
| **Putative excitatory** | (82.44%) 205 | (67.15%) 164 | (52.58%) 91 | (52.58%) 84 |
| **Putative inhibitory** | (17.56%) 17 | (32.85%) 21 | (47.41%) 22 | (47.41%) 16 |
| **Total** | **222** | **185** | **113** | **100** |

A between-group difference was detected, however, when we compared the amplitudes of taste responses. Figure 3E shows the average responses to sucrose (left) and CA (right) in apoE4 and WT animals, before and after CTA. Note that the FRs of each neuron were normalized to its BL to eliminate between-group differences in BL activity (Fig. 3B). These graphs clearly show that apoE4 GC taste responses (in red) are weaker than those of the WT group (in green); independent statistical analyses of sucrose (t(618)=4.29, p<0.0001) and CA responses (t(618)=4.45, p<0.0001) confirmed this observation (Fig. 3F). Furthermore, this analysis revealed apoE4 conditioned stimulus (sucrose) response plasticity deficits: as expected, neuronal responses to the non-conditioned taste CA remained unchanged following CTA learning for both genotypes (Tukey: WT Pre vs. Post p=0.82, apoE4 Pre vs. Post p=0.9), and responses to the conditioned sucrose changed in the WT group (Tukey: WT Pre vs. Post p=0.02); but were not affected by training in the apoE4 group (Tukey: apoE4 Pre vs. Post p=0.9). Note that the CTA-related changes observed in control rats appear, as expected, to be mainly confined to later aspects of the responses: GC single-neuron responses encode taste-specific information in a sequence of epochs in the 1-2 sec following taste stimulation—an early epoch (EE) that codes taste identity, and a late epoch (LE) that codes palatability (Katz et al., 2001; Piette et al., 2012). CTA is associated with activity changes primarily in the palatability-rich LE (Grossman et al., 2008; Moran and Katz, 2014). It appears that our current data replicate this result, and suggest that this LE response plasticity fails to develop in apoE4 rats. To test this observation, we measured the response amplitudes for sucrose in these epochs and tested for differences between the groups across days. Our results show that the overall higher response FR in the WT group compared to apoE4 was apparent in both the EE and LE epochs (Fig. 3G. ANOVA, genotype: EE: F(1,616)=11.69, p<0.001; LE: F(1,616)=21.89, p<0.0001). During the EE, neither genotype showed learning-related modifications (Fig. 3G, Tukey: WT Pre vs. Post p=0.62, apoE4 Pre vs. Post p=0.81). Learning effects appeared in the LE, where as expected, post-CTA responses increased in WT animals (Fig. 3G, Tukey: WT Pre vs. Post p<0.001). In the apoE4 animals, on the other hand, the FR in the LE remained low and similar to the apoE4 pre-CTA value (Fig. 3G, Tukey: apoE4 pre vs. post: p=0.9). Analysis of CA taste responses, which rats were not trained to avoid, showed only a genotype effect (Fig. 3H, genotype: F(1,616)=20.42, p<0.0001; day: F(1,616)=0.12, p=0.72; Interaction: F(1,1)=0.02, p=0.87). Together, these results show that apoE4 is correlated with lower BL and evoked responses in GC neurons, as well as with insufficient post-CTA modulation in GC-LE taste responses, a specific plasticity-driven process that underlies CTA learning.

The FR anomalies revealed in apoE4 animals could be the results of cell-specific alterations, originating from either excitatory or inhibitory neurons. Previous research suggested that inhibitory neurons are more sensitive to the apoE4 induced neurotoxicity showing higher level of neuronal loss compared to pyramidal neurons (Leung et al., 2012; Liraz et al., 2013; Tong et al., 2014; Nuriel et al., 2017; Najm et al., 2019). Therefore, the FR anomalies revealed in apoE4 animals could manifest differently in different cell types. To address this possibility, we used a spike-shape classification method that relied on the ratio between the height and width (H/W) of the neuron’s averaged action potential to separate the neurons into either putative inhibitory or excitatory. The distribution of the H/W ratio across all recorded neurons showed a clear bi-modal pattern (Fig. 4A). Based on previous studies we defined neurons from the high (red) and low (blue) H/W groups as putative inhibitory and excitatory neurons, respectively (Fig. 4A, Yokota et al., 2011; Samuelsen et al., 2012; Samuelsen and Fontanini, 2017). The number of neurons collected from each experimental group is summarized in Table 1.

**Figure 4.**
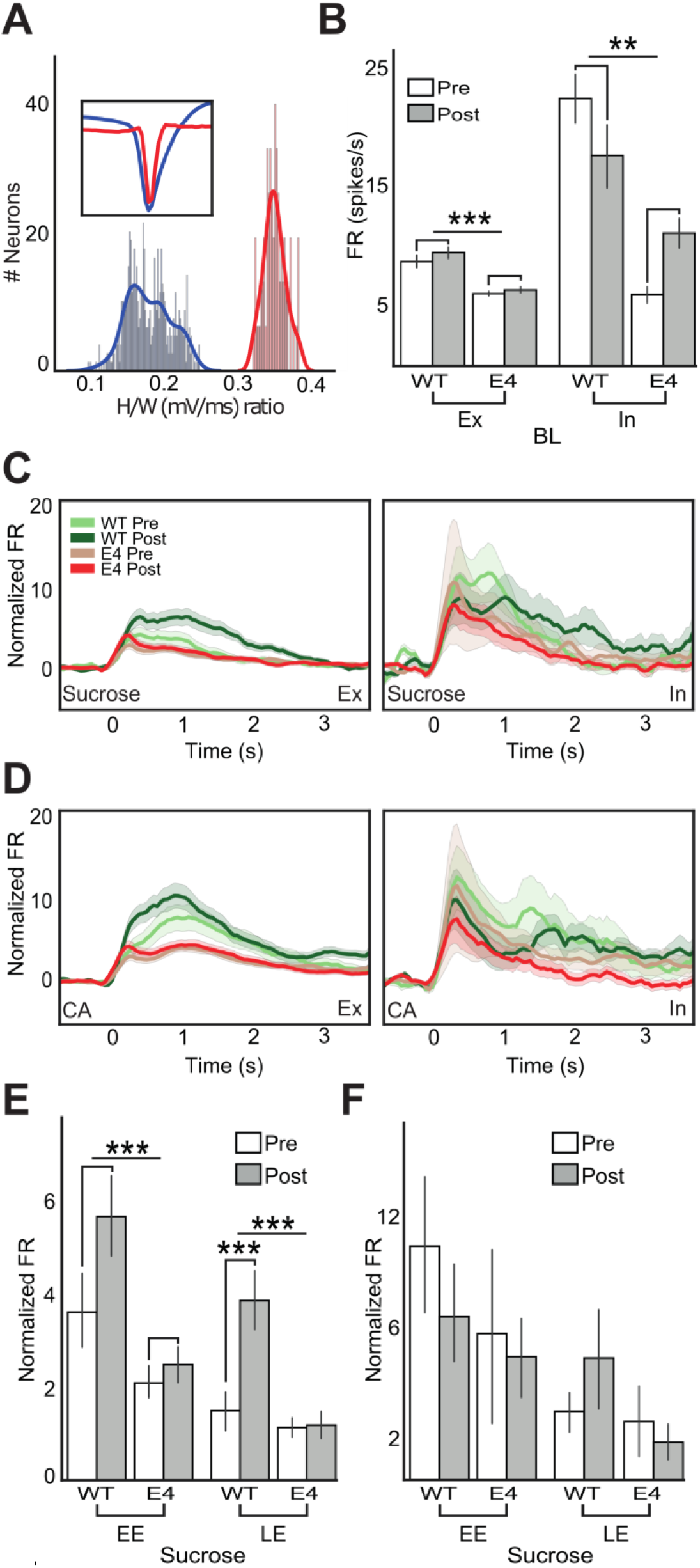
The impact of apoE4 on excitatory and inhibitory neurons. **A)** A histogram of the ratio between the height (H) and width (W) of each neuron’s spike shape shows a clear bi-phasic distribution. Inset represents the averaged normalized waveform of the two clusters, which were interpreted as inhibitory (red) and excitatory (blue) neurons. **B)** Mean BL activity revealed a genotype effect for both excitatory and inhibitory neurons. **C)** The averaged response to sucrose of excitatory (left) and inhibitory (right) neurons over 3 seconds after taste delivery (at 0), normalized to the BL activity. **D)** As in (C) but for CA. **E)** Excitatory neurons’ averaged responses to sucrose in the early (EE) and late (LE), before (Pre) and after (Post) CTA, normalized to BL activity. Apoe4 induced lower FR in both response epochs and prevented post-CTA LE FR increase. Two-way ANOVA: EE: day: F(1,540)= 2.88, p=0.08; genotype: F(1,540)=15.86, p<0.0001; Interaction: F(1,1)=1.97, p=0.16. LE: day: F(1,540)=4.97, p=0.026; genotype: F(1,540)=15.52, p<0.0001; Interaction: F(1,1)= 9.06, p=0.002 **F)** Inhibitory neurons’ averaged responses to sucrose. Both EE and LE showed no significant effect. Two-way ANOVA: EE: day: F(1,72)=0.53, p=0.46; genotype: F(1,72)=0.93, p=0.33; Interaction: F(1,1)=0.13, p=0.71; LE: day: F(1,72)=0.19, p=0.65; genotype: F(1,72)= 2.03, p=0.15; Interaction: F(1,1)=1.01, p=0.31. In = inhibitory neurons; Ex = Excitatory neurons; EE = 0-1 second post taste delivery; LE = 1-2 second post taste delivery; Normalized FR = the FR of each neuron normalized to its BL activity; **p<0.01, ***p<0.001

Cortical inhibitory neurons have higher BL FR than excitatory neurons (Yokota et al., 2011). Correspondingly, we found that in WT rats the BL FR of inhibitory neurons was significantly higher than the excitatory neurons (Fig. 4B), further validating our classification method. Overall, our analysis revealed that apoE4 reduces BL activity of both cell types (Fig. 4B, Two-way ANOVA; Excitatory: genotype: F(1,540)=17.39, p<0.0001, interaction: F(1,1)=0.3, p=0.58; Inhibitory: genotype: F(1,72)=7.94, p=0.006; Interaction: F(1,1)=1.89, p=0.17). The BL-FR of inhibitory neurons in the apoE4 group was exceptionally low, reaching about a quarter of the WT group before CTA. As expected, no learning-related changes were found in the averaged BL of the two cell type in both groups (Fig. 4B). We further tested whether theapoE4 reduced BL excitability has a cell-specific influence over taste responses. The averaged response magnitude to sucrose (Fig. 4C) and CA (Fig. 4D) was computed separately for excitatory and inhibitory neurons (left and right panels, respectively) across groups and days. These figures show that apoE4 seems to attenuate taste responses in both cell types, but with a stronger effect in excitatory neurons. Epoch-specific analyses of sucrose responses confirmed that apoE4 significantly reduces taste responses of both epochs, and prevents the post-CTA LE FR increase observed in the WT group (Fig. 4E). The group differences observed in excitatory neurons did not replicate in the inhibitory neurons (Fig. 4F). These results suggest that the impaired capacity of young animals’ GC neurons to undergo plasticity modification following CTA (Fig. 3H) stem mainly from deficits in excitatory neurons.

So far we have shown that cortical neurons in young adult apoE4 rats are less active, respond weakly to tastes, and fail to increase their response to the conditioned taste to support CTA learning. At least two hypotheses could be proposed to explain the relationship between the apoE4 electrophsiological abnormalities and impaired learning. Perhaps the low FR of apoE4 neurons impairs neuronal coding of taste information, thereby interfering with taste perception and consequently with learning. It is also possible that, despite the lower FR, taste information is largely intact, and additional impairments in the ability to undergo neuronal plasticity is the cause of learning difficulties.

To test these hypotheses, we compared the encoding of taste information in WT and apoE4 GC neurons at the single neuron and population levels. Observing apoE4-related interference with taste coding prior to learning will suggest an innate taste perception malfunction, while intact taste coding but lack of coding plasticity following CTA will support plasticity deficits. At the single neuron level, taste information exists in the response FR if at least one taste response is different from the others (Katz et al., 2001; Piette et al., 2012, see the Methods section). Figure 5A shows two representative GC neurons—one that carries taste-specific information (Fig. 5A left), and one that does not (Fig. 5A right). Comparing the number of taste-specific neurons across the different genotypes and conditions did not reveal significant differences (χ^2^ test, χ^2^(9,620)=6.32, p=0.99, see Table 1). Furthermore, responses in WT and apoE4 rats were similarly capable of correctly identifying the identity of the taste that evoked the neural response (χ^2^ test, χ^2^ (3,620)= 0.59, p=0.09,Fig. 5B-E).

**Figure 5.**
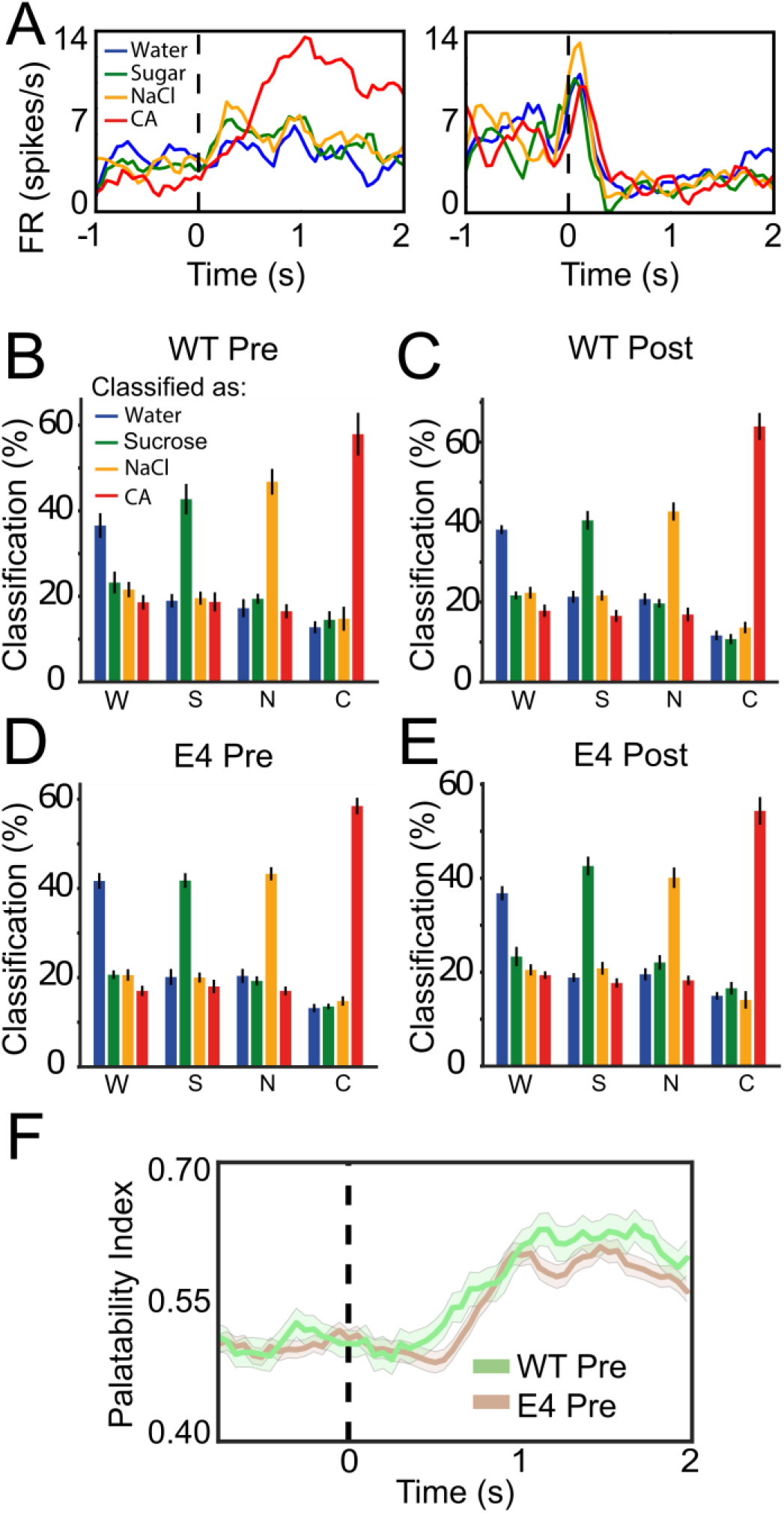
GC neurons of apoE4 and WT animals code similar taste information. **A)** Left panel: a taste-specific neuron; Right Panel: a non-taste specific neuron. **B-E)** A single neuron, response-pattern based classification revealed similar successful classification of all tastants across all groups **F)** GC single neuron palatability information coding shows similar increase in the LE in both genotypes before CTA. Classification = the percentage by which a single neuron taste stimulus response was classified as evoked by a certain tastant; W=Water; S=Sucrose; N=NaCl; C=CA; WT = Wildtype; E4 = apoE4; Pre = before CTA.

We also compared the intensity of palatability-relatedness of LE responses (Katz et al., 2001; Piette et al., 2012; Sadacca et al., 2012) in WT and apoE4 rat single neurons. We defined a palatability index (PI) based on the correlations between the taste responses and their known innate palatability (Sadacca et al., 2012; Moran and Katz, 2014); the fact that this index increased in the LE of apoE4 GC neurons similar to the WT group (Fig. 5F) suggests intact palatability processing in the apoE4 animals. It therefore appears that, despite apparent differences between the FR of WT and apoE4 GC neurons, apoE4 and WT animals show similar taste identity and palatability encoding capacities.

Of course, GC neuronal coding, and specifically the activity changes that underlie CTA learning, are best described in the neuronal ensemble level (Jones et al., 2007; Moran and Katz, 2014). We therefore asked whether the reduced activity of the apoE4 GC neurons may affect the network’s palatability coding (PC) space, and its relation to the rats’ impaired learning. To that end, we considered the response of the ensemble to a taste as a trajectory in N-dimensional space (N denotes the number of neurons in the ensemble) (Samuelsen et al., 2012). Examples of these trajectories (reduced to 2D space) are presented in figure 6A. Each of the colored trajectory lines represents the response of an ensemble to one of the 4 tastes across 3 seconds in a single rat. The trajectories for different tastes typically begin similarly, and then diverge in the late parts of the responses.

**Figure 6.**
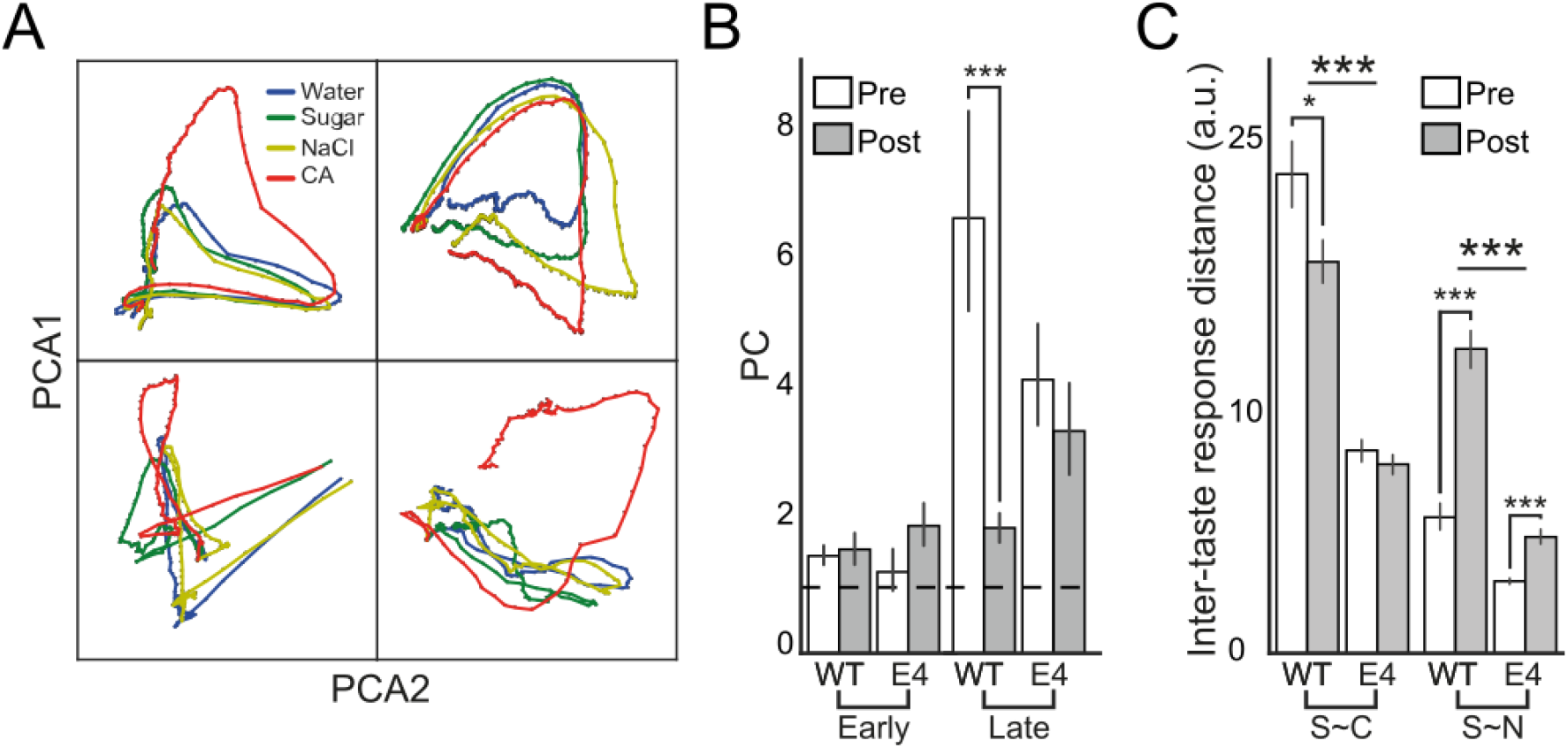
ApoE4 does not attenuate innate population PC but interfere with the PC update required for CTA. **A)** Four examples of GC ensemble response dynamics over the 3 seconds post-taste delivery projected on a 2D space. i) Similar dynamics for all tastant except for CA; ii) Initial uniform dynamics that later separate: iii) A dynamic that scatters without clear distinction between tastants; iv) A network that can only clearly separate the response for CA towards the end. **B)** Sucrose PC across genotypes, epochs and days of experiment. We found similar low PC values across genotypes and days during the EE. In the LE, the WT group showed high PC before CTA that was decreased following CTA, while in the apoE4 group sucrose PC was not affected by learning. **C)** The LE distance ratio separated into its components shows lower S~C and S~N in the E4 compared to WT. CTA affected a specific decrease in S~C distances only in the WT animals and an S~N increase in both genotypes which was more prominent in the WT group. PC = Sucrose palatability coding; Early = 0-1 second post sucrose delivery; Late = 1-2 second post sucrose delivery; Pre = before CTA learning during the training session; Post = after CTA learning during the test session; WT = Wildtype; E4 = apoE4; ***p<0.001

We assessed the ensemble sucrose PC (Moran and Katz, 2014)by calculating the ratio between the distances between the sucrose trajectory and those in response to CA and NaCl (S~C and S~N respectively). The higher the value of PC (sucrose ensemble trajectory is far from the aversive CA trajectory and close to the palatable NaCl), the more sucrose is coded as palatable. As expected, PC values were low and similar across groups and days during the EE, which does not code for palatability (Fig. 6B. Two-way ANOVA; day: F(1,276)=3.29, p=0.07; genotype: F(1,276)=0.77, p=0.38; interaction: F(1,1)=0.05, p=0.82). In the LE, meanwhile, sucrose was coded as highly palatable by the population activity of both genotypes prior to CTA (i.e., high PCs, albeit lower for apoE4; Tukey, WT pre vs. E4 pre, p<0.001) (Fig. 6B). Between-group differences were found following CTA: while the sucrose aversion caused a significant reduction of WT PC (Tukey, WT pre vs. WT post, p<0.001), PC remained unchanged in the apoE4 group (Tukey, E4 pre vs. E4 post, p=0.46). Inspection of the two PC components found that the lower FR in the apoE4 group caused a reduced ensemble PC space (Fig. 6C); Pre-CTA, S~C and S~N distances were significantly reduced in apoE4 animals to begin with (Tukey: S~C: E4 pre vs. WT pre p<0.001; S~N: E4 pre vs. WT pre p<0.001).

In WT rats, the reduction of sucrose palatability following CTA was the combined result of a decrease in the distance from aversive CA (Lower S~C, Tukey WT pre vs. post: p=0.016) and an increase in the distance from NaCl (higher S~N, Tukey WT pre vs. post p<0.001). In the apoE4 animals the S~N distance showed a much smaller increase (Tukey E4 pre vs. E4 post: p=0.001), and S~C distance failed to change post-CTA at all. Together, these results suggest that although sucrose identity and palatability could be sufficiently coded by GC neurons of the apoE4 animals (despite general reductions in FR), these responses fail to allow sufficient coding space for plasticity to support CTA learning.

## Discussion

AD is believed to start decades before the appearance of its characterizing pathologies (Khan, 2018; Younes et al., 2019). Impaired basal synaptic plasticity was suggested to underlie these early apoE4 dysfunctions and the subsequent development of brain pathologies in apoE4 carriers (Teter, 2004; Huang, 2011; Verghese et al., 2013; Matura et al., 2014; Michaelson, 2014; Abushakra et al., 2016). Our recent publication supports this hypothesis by showing plasticity deficits in extra-hippocampal brain regions of young TR apoE4 mice which were correlated with impaired CTA learning (Har-Paz et al., 2019). We further hypothesized that these early deficits should impact neuronal activity and coding that could explain the impaired CTA learning. Therefore, we recorded neuronal activity from the GC of hApoE4 and WT SD rats before and after CTA learning. Behaviorally, the young hApoE4 rats replicated our previous results in mice showing impaired CTA acquisition (Fig. 2B), despite intact taste perception (Fig. 2C-E), and taste coding (Fig. 5) prior to learning. Electrophysiogically, we observed reduced apoE4 GC neuronal activity at BL and evoked responses compared to WT, in both inhibitory and excitatory neurons (Fig. 3, 4). In line with the behavioral results, taste responses in the GC adequately increased after CTA in the WT group but did not change in apoE4 animals (Fig. 3). Additionally, we show that the GC neurons of the apoE4 group failed to appropriately update their sucrose palatability coding in the LE following CTA training, an update that is crucial for CTA (Fig. 6). Our combined results from apoE4 mice and rats show that apoE4 reduces both structural and functional plasticity modalities that impact basal and taste-evoked activity of young brains that interrupts with normal learning processes at young ages, before the appearance of the late AD pathologies.

Contrary to the results shown in the current study, the majority of the apoE4 neuronal activity in mice (Klein et al., 2014; Tong et al., 2016; Nuriel et al., 2017) and human (Bookheimer et al., 2000; Trivedi et al., 2008; Filippini et al., 2009) studies, reported an apoE4 neuronal hyper-activation. Those findings, however, are reported from research conducted on aged subjects in the hippocampus and therefore may be influenced by advanced degeneration processes (Andrews-Zwilling et al., 2010; Chen et al., 2010; Mannix et al., 2011; Yin et al., 2011; Matura et al., 2014; Kim et al., 2018). This body of evidence attributes the observed hyper-activation to the increased vulnerability of hippocampal γ-aminobutyric acid (GABA)-expressing interneurons to the neurotoxicity induced by the apoE4 genotype. This may lead to GABAergic loss resulting in excitotoxicity and an overstimulation of glutamatergic receptors, which can inflict neuronal degeneration associated with AD (Leung et al., 2012; Belsky et al., 2013; Nuriel et al., 2017; Najm et al., 2019). It is therefore conceivable that the observed early cortical hypo-activity originates from early impaired plasticity mechanisms (Teter, 2004; Wang et al., 2005; Har-Paz et al., 2019), rather than apoE4-induced neurotoxicity that plays a role at later ages. If so, this interesting transition between hypo- to hyper-activation may reflect the progression of more pathological effects of the apoE4 phenotype.

The question then remains, what is the mechanism that links the apoE4-related decreased structural plasticity in the GC to their FR and learning aberrations. A possible answer is that apoE4 reduces the ability to create novel GABAergic and glutamatergic vesicles, whether basally or following taste stimuli, that consequently limits evoked spiking activity by preventing fast replenishment of release-ready vesicles (Wesseling and Lo, 2002; Chen et al., 2010; Har-Paz et al., 2019). The disruption in GABAergic or glutamatergic functions may impair the formation of novel-taste memory and CTA (Tucci et al., 1998; Miranda et al., 2002; Daniel et al., 2016; Sandoval-Sánchez et al., 2020). This hypothesis is directly supported by our previous study showing CTA-induced vesicle reserves deficits in the GC of apoE4 mice (Har-Paz et al., 2019). We therefore anticipated that apoE4 anomalies will mostly manifest in evoked post-CTA FRs.

Surprisingly, we found that before CTA the apoE4 basal FR averaged to about half of that of the WT group across the entire population, an effect that was more pronounced in inhibitory neurons. The fact that our previous study found similar basal levels of inhibitory vesicles in the GC suggests that the lower basal FR of inhibitory neurons may be a lack of sufficient network net excitation rather than insufficient cellular resources (Stone et al., 2011; Har-Paz et al., 2019). However it is also possible that local synaptic interactions within the GC such as apoE4 reduced receptor signaling can account for the observed phenotype (Chen et al., 2010). With regard to taste responses, the low apoE4 evoked potentials is likely a combination of the above-mentioned local cellular structural plasticity anomalies and similarly reduced input coming from connected neurons. Importantly, the diminished apoE4 responses are not a consequence of the low basal FR as we normalized each neuron response to its BL. The systemic malfunction in driving spiking activity can cause an accumulative effect which can be detected already at BL but becomes more apparent as the neuron is required to increase its response to stimuli. The apoE4-related reduction in evoked responses was only noticeable in excitatory neurons, however this may be due to high inhibitory FR variability and their smaller sample size. Regardless, This FR rigidity can account for the observed lack of apoE4 LE changes as a response to the conditioned tastant following CTA (Fig. 3H, Grossman et al., 2008; Moran and Katz, 2014)

GC neurons uniquely code information in their dynamic spiking activity (Katz et al., 2001; Jones et al., 2007). Theoretically, the overall reduced apoE4 GC FR (Fig. 3, 4) may affect their neural network’s information capacity (Stevens and Zador, 1996; Schneidman et al., 2000), attenuate the ability to differentiate between stimuli in short time scales, thus interfere with learning. It is possible, then, that the observed learning impairments are the results of sensory misperception rather than deficits in neuronal plasticity. Our behavioral test, however, found similar sucrose preference between apoE4 and WT rats (Fig. 2E). In congruent with the behavioral tests, analyses of taste identity and PC performance by GC neurons, both in the single neuron (Katz et al., 2001; Piette et al., 2012; Sadacca et al., 2012) and the population (Jones et al., 2007; Moran and Katz, 2014; Mukherjee et al., 2019) levels, found no differences between the genotypes (Fig. 5). However, both excitatory and inhibitory neurons did not manage to change their responses to support CTA (Fig. 4D, E), which disrupted the update of sucrose population PC (Fig. 6C). Collectively, these results support the hypothesis regarding an overall plasticity dysfunction in young apoE4 animals’ single neurons and networks that may impact their ability to learn.

While these results are consistent with our previous study in mice and further characterize apoE4 plasticity deficits, they are not without their limitations. The most notable one is the lack of humanized apoE3 matching control rats. Current apoE4 research mostly focuses on the TR mice models, but our electrophysiological study required larger brains and therefore we chose to utilize the recently-developed hApoE4 rats. This decision, however, prevented the usage of matching hApoE3 control groups, as this genotype has not yet been developed. Instead we used WT SD rats, the background strain for the hApoE4 rats. We argue, however, that SD rats can still serve as valid controls in our study. First, the behavioral results have been replicated between apoE4 expressing rats and mice and TR-apoE3 control mice showing striking similarity in their behavior following CTA to their wildtype littermates and WT SD rats. Specifically, both genotypes showed pronounced aversion to weak and strong CTA. Moreover, apoE3 and WT mice displayed similar extinction rate following repeated exposures to the CS taste without the LiCl reinforcement (data not shown). While we cannot refute the possibility that our results are a mere consequence of the genetic manipulation, our data suggest it is highly unlikely.

Overall, our findings hold a great importance to the apoE4 field by exposing the early, possibly innate, apoE4 impact on brain activity and functional plasticity. Our results call for a careful evaluation of the apoE4-related neural molecular deficits alongside their functional electrophysiological correlates, as these two modalities are likely to co-influence each other. Moreover, our results highlight the impact of apoE4 on extra-hippocampal regions, and its influence on the functional plasticity domain. These early behavioral, molecular and functional deficits are likely to play a role in the development of later apoE4 related pathologies. This may allow these phenotypes to act as markers for apoE4 early clinical pathology detection, as well as potential targets for early interventions (e.g. upregulate plasticity) to slow down or even prevent the development of neurological conditions like AD.

## Notes

### Competing Interest Statement

The authors have declared no competing interest.

